# Patterns and drivers of genome-wide codon usage bias in the fungal order Sordariales

**DOI:** 10.1101/2025.05.07.652452

**Authors:** Noah Hensen, Markus Hiltunen Thorén, Hanna Johannesson

## Abstract

Here we present a study on amino acid composition, codon usage bias (CUB), and levels of selection driving codon usage in Sordariales fungi. We found that GC ending codons are used more often than AT ending codons in all Sordariales genomes, but the strength of CUB differs amongst families. The families *Podosporaceae* and *Sordariaceae* contain relatively low genome-wide levels of CUB, while the highest levels of CUB are found in *Chaetomiaceae* and the “BLLNS”-group, a monophyletic group of five other Sordariales families. Based on genomic clustering, and ancestral state reconstruction of GC nucleotides at the third codon position, we hypothesize that *Podosporaceae* and *Sordariaceae* represent the ancestral state of amino acid composition and CUB. The *Chaetomiaceae* and BLLNS have most likely diverged from this state, with increased natural selection driving use of specific codons, resulting in higher genome-wide CUB. We expect that the higher levels of CUB in *Chaetomiaceae* genomes might have been caused by ecological niche specialization, including high optimal growth temperature of some *Chaetomiaceae* species.

## 1. Introduction

In messenger RNA, triplets of nucleotides are read simultaneously during translation. These so-called codons form a unit of genomic information encoding a particular amino acid. In the standard genetic codon table, a total of twenty amino acids is encoded by 61 codons, with three additional stop-codons signaling termination of protein synthesis. The same amino acid can thus be encoded by different codons (synonymous codons). Both methionine and tryptophan are encoded by one codon each, while the other eighteen amino acids are encoded by multiple synonymous codons (1–3).

The frequency with which certain amino acids are used to build proteins is well-conserved across species, and even across kingdoms (4). Yet, deviations from the average composition exist, and have been linked to, e.g., cellular organization, gene expressivity and enhancement of protein stability in response to environmental pressures, such as high ambient temperatures (5,6). Synonymous codons to encode for these amino acids are not randomly or equally used, however. Instead, some codons are consistently used more often than others. The phenomenon of using synonymous codons with different frequencies is termed codon bias or codon usage bias (CUB) (3). It influences diverse cellular processes, such as RNA processing, protein translation and protein folding. The level of CUB is furthermore a critical factor in determining gene expression and cellular function (3).

Codon usage bias is a well-established phenomenon (7). It is found in all organisms, but there is a large variation in which codons are preferentially used for a particular amino acid. The level of bias also varies amongst species, and amongst genes within an organism’s genome (3,8). These differences in the level of CUB can arise from multiple factors, such as mutational biases, selection for translation efficiency, and genetic drift (3,9,10). It is generally assumed that codon bias reflects a mutation-selection balance (3,11). Analyzing CUB of closely related species thus helps to identify the main forces that drive their evolution, and is an important step in evolutionary studies.

Within the fungal kingdom, the order Sordariales is a large and diverse group of fungi with high scientific importance (12–15). The order includes species used as model-organisms for basic cellular processes, such as *Neurospora crassa* and *Podospora anserina* (12–15), as well as species that produce a wide range of biologically active secondary metabolites (16,17). Within the Sordariales, the *Chaetomiaceae* family is well-known for containing multiple thermophiles (18,19), a trait correlated with multiple genomic traits (18,20). Recent research by Steindorf et al., (2024), for example, showed a correlation between optimal growth temperature in fungi and variation in the number of codons used in protein coding genes, as well as GC level at the third codon position (18). However, a broad exploration of the CUB trends in the *Chaetomiaceae* family, or the rest of the Sordariales order, has not yet been presented.

In this study we describe the amino acid composition and codon usage variation in Sordariales by analyzing whole-genome sequences from 99 species belonging to nine families in the order (21, and references therein). We identify patterns of codon usage across the order, and analyze the signatures of selection driving CUB across different families (21). Our study contributes to the understanding of fungal genome evolution in the Sordariales, and is valuable for inferring evolutionary trajectories of genomes and protein-coding genes. Furthermore, analyses of amino acid composition and CUB has potential future biotechnological application in increasing heterologous expression of important secondary metabolites (3,16,22).

## 2. Methods

As outlined in detail below, we analyzed a range of different indices to describe amino acid composition and codon usage across the Sordariales order, and the processes influencing them. The relative synonymous codon usage and the effective number of codons were used to assess overall strength of codon bias across the order (3,23,24). The codon adaptation index was used to calculate CUB of highly expressed genes versus the rest of the genome (25,26), and we contrasted selection versus mutational pressure on codon usage with neutrality plots and assessed the effective number of codons in relation to GC content at the third codon position (24,27).

### Sequence data collection and filtering

In our study, we make use of a large dataset of genomic information and a well-supported phylogeny of the fungal order Sordariales that have been made available recently by Hensen, et al., 2023 (21 and references therein). The dataset contains genomes of species from nine Sordariales families, of which three (*Sordariaceae, Podosporaceae* and *Chaetomiaceae*) contain a large enough number of genomes to make comparisons of intra- and interfamily variation. Additionally, five of the smaller families (*Bombardiaceae, Lasiospaeriaceae, Lasioshaeridaceae, Naviculisporaceae*, and *Schizotheciaceae*) form a strongly supported monophyletic group in the Sordariales phylogeny (21). They were here combined into one group for statistical analysis. From here on out, we refer to this group of families as the “BLLNS” group. The family *Diplogelasinosporaceae* is only represented by one genome, which groups independently from the BLLNS. As such, we do not include this species in statistical comparisons. The outgroup was represented by three Sordariomycete genomes from outside of Sordariales (*Eutypa lata, Lollipopaia minuta, Phialemonium atrogriseum*)(21)(Table S1).

High-quality coding sequences (CDS) were extracted from the annotated genomes and used in downstream analysis. These high-quality genes were selected based on the following criteria: the spliced CDS 1) have a length of ≥ 100 bp, 2) contain no partial codons, 3) have no internal stop codons, and 4) contain both start and stop codons (Table S2). Only the coding sequences of the remaining genes were used for further analysis.

### Patterns of amino acid usage and codon usage

The frequency of amino acids was estimated for each genome, and patterns of codon usage were analyzed with relative synonymous codon usage (RSCU) values. The RSCU values give the observed frequency of a codon divided by the expected frequency under the assumption of equal usage of synonymous codons (3). Codons with an RSCU value of 1 are regarded as unbiased. Values > 1 indicate that there is a higher frequency of a particular codon in the genome than expected under random use, and a value < 1 indicates that a codon is found less frequently than under random use. Codons with RSCU values > 1.6 and < 0.6 are considered as “overrepresented” and “underrepresented” codons, respectively (see e.g. 28). The average frequency of GC nucleotides at the third codon position (GC3) was used as an additional index for codon bias (3,29–31). The average amino acid composition, RSCU frequencies, and the GC3 were calculated on genome-wide CDS with BioKIT V0.1.2 (32).

To systematically investigate CUB variation across different families of the Sordariales, we performed automatic clustering of the genomes for both amino acid composition and RSCU. Automatic clustering was performed with the pheatmap R-package (33). We additionally obtained a kingdom-wide phylogeny and GC3 contents for 689 fungal species from Wint et al (2022) (34). Ancestral state reconstruction of GC3 contents was performed as described by Hensen et al., (2023), using the R package Phytools V0.6.44 with function ContMap (21,35).

### Estimating genome-wide strength of codon usage bias

We analyzed the genome-wide codon usage bias by calculating the effective number of codons (ENC) and the codon adaptation index (CAI). The ENC indicates the magnitude of codon bias by calculating how many of the 61 possible codons are used in a gene (24). ENC values start from 20, indicating one codon was exclusively used to code for a given amino acid, and range up to 61, indicating all codons were used equally (24). An ENC value ≤ 35 suggests that a gene possesses a strong codon bias (3).

The codon adaptation index (CAI) is based on the assumption that CUB is optimized for efficient translation of highly expressed genes, and is assessed by comparing the codon usage of a gene to the codon usage of highly expressed genes (36). Genes encoding ribosomal proteins are generally highly expressed (36) and were used as the reference set against which CAI values were calculated. To identify ribosomal genes, we translated the genes of each genome with seqkit translate (37,38). We next ran interproscan (39,40) on the set of translated genes with the -appl Pfam option and compiled a list of ribosomal genes from the PFAM eukaryotic ribosomal genes PFAM database (Table S3). The genes matching the set of PFAM ribosomal genes were isolated from each genome. CAI values lie between 0 and 1, where a lower CAI implies that preferred codons are assigned only to a small group of highly expressed genes (26). Values above 0.59 indicate no differential use of synonymous codons between highly expressed genes and the rest of the genes in the genome (26), and further suggest that a genome has experienced stronger genome-wide selection to maintain a specific CUB that is optimized for efficient translation (25). The software CodonW V1.4.2 was used to run a correspondence analysis on the ribosomal genes, which was then used as a reference input for CAI. The ENC and CAI were calculated on genome-wide CDS, using CodonW V1.4.2 with the - totals setting (41).

### Disentangling selection from mutational bias as factors driving codon usage bias

Neutrality plots and ENC/GC3 plots were used to infer whether codon usage can be attributed to selection or mutational bias. The analyses were done both on the entire set of coding sequences and on single genes. Neutrality plots were created by plotting the average GC at the first and second codon position (GC12) on the Y-axis, versus GC3 on the X-axis. Analyses of selection by the use of neutrality plots assume the third codon position is neutral (27,42). Under complete neutrality, all positions in the codon are equally likely to mutate, which leads to a statistically significant correlation between the GC12 and the GC3 content, and a slope of the regression line close to 1. Selection driving codon usage, on the other hand, can lead to a slope close to zero or an overall lack of correlation between GC12 and GC3 (43–45). In neutrality plots with significant correlations, the absolute slope value times 100 is equal to the percentage of mutation bias (27,42,46). The level of selection is then described as

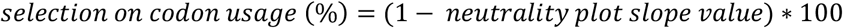

Neutrality plots were created for each genome individually, with each individual gene represented by a discrete point. To draw neutrality plots, the GC levels of the first, second and third codon position were analyzed for each gene using BioKIT V0.1.2 (32). Regression lines of neutrality plots were determined using the R-package ggpubr (47,48), and absolute slope values were used to estimate genome-wide levels of selection versus neutrality. Additional to neutrality plots, ENC/GC3 plots were used to determine whether the codon usage of single genes is affected mainly by mutation or mainly by selection. A normal distribution curve represents the expected ENC values plotted against expected GC3 values. When the usage of codons is limited only by G + C mutation bias, the genes represented by points in the ENC-GC3s plot are placed on this line. The expected ENC values were calculated according to the following function:

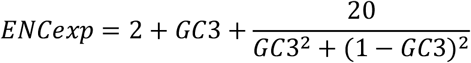

Alternatively, factors such as natural selection lead to a relative decrease in GC3 content compared to the ENC, causing genes to be distributed below the curve (Wright, 1990). Genes (*dNC*_*sg*_) where ENCobs < ENCexp (*dNC*_*sg*_ < 0) were taken as genes that displayed signs of selection driving codon usage.

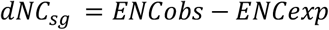

The percentage of genes under selection for codon usage (*dNC*) was used to estimate distribution of selection over the genomes.

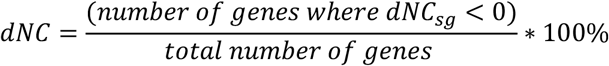

Both the slope of neutrality plots and percentage of genes showing signs of natural selection were based on individual, high-quality genes. Neutrality plots and ENC/GC3 plots were drawn with R-package ggplot2 (48).

### Statistical analysis

Tests of normality were performed for each genome property (ENC, CAI, GC3, dNC, and absolute slope of neutrality plots). In all cases, Shapiro-Wilk normality tests showed p < 0.05; indicating that normality cannot be assumed. Statistical comparisons of trait values amongst the families were done with Pairwise Wilcoxon Rank Sum Tests with the R function pairwise.wilcox.test with a Bonferroni p-adjustment method.

Pearson’s correlation coefficient was additionally used to test for correlations among the trait variables on the entire phylogeny without outgroups (R-function cor.test, with method = “pearson”). To correct for phylogenetic dependence of species traits, the R package ape was used to compute phylogenetically independent contrasts (49,50).

### Figures and scripts

All plots and statistics in this manuscript were obtained with R V4.4.1 in Rstudio (51,52). Our developed procedures and analyses consist of custom bash and R scripts, which we have made available together with instructions for use and a list of used software versions and R-packages, at https://github.com/NoahH35/CodonUsageBias.

## 3. Results and discussion

### 3.1 Amino acid frequencies are conserved across the fungal order Sordariales

The amino acid composition has been described to remain largely similar across species and kingdoms (5,53,54). In fungi, plants and prokaryotes, research showed that -of the 20 standard amino acids-alanine, leucine and serine were present in the highest frequencies, while cysteine, histidine, methionine, and tryptophan frequencies were the lowest (5,53,53). The amino acid composition in the Sordariales was in line with these general findings (figure 1). However, just as genomic traits such as GC content, genome size, and gene number differ amongst the Sordariales families (21), small differences in amino acid composition were found (figure 1, table S4). The *Podosporaceae* and *Sordariaceae* showed a relatively more even frequency of amino acid usage, while the amino acid frequencies in *Chaetomiaceae* and the BLLNS showed slightly higher bias towards the use of particular amino acids. Specifically, if all 20 standard amino acids would be used equally, the frequency of all would be 5%. Instead, alanine, for example, represented 9.6% of the total amino acids in *Chaetomiaceae*, and 9.2% in the BLLNS, versus 8.7% and 8.5% in *Sordariaceae* and *Podosporaceae* respectively.

**Figure 1.**
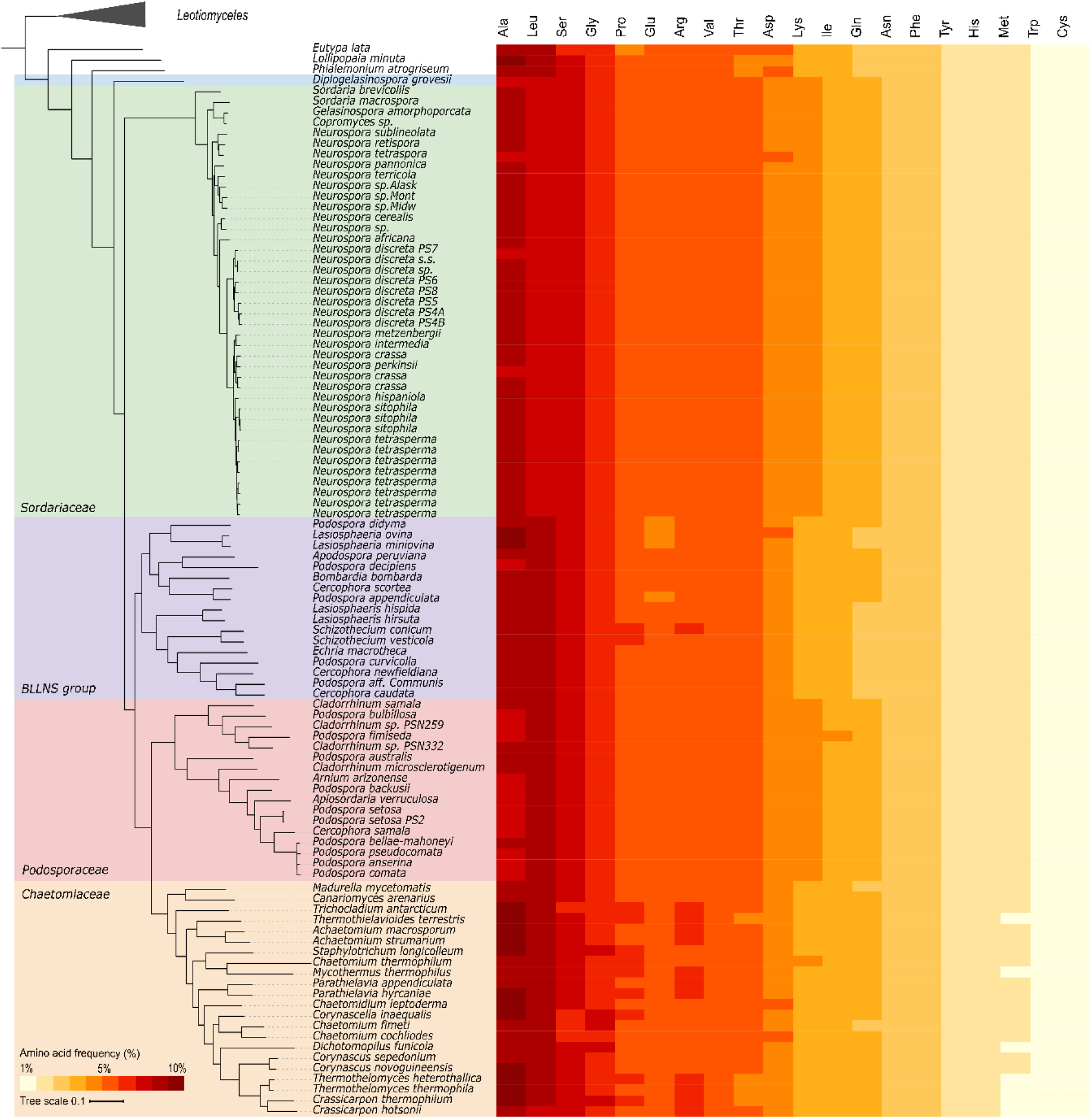
Amino acid composition across the Sordariales. Average amino acid frequencies (%) for each of the investigated genomes. Amino acids are ordered from most used (left, dark-red) to least used (right, light-yellow) for the average of all genomes combined.

### 3.2 G and C ending codons are overrepresented in the Sordariales order

All Sordariales families showed similar trends for which codons were preferentially used to encode a particular amino acid (figure 2). The RSCU and GC3 analysis showed that GC ending codons were used more often than AT ending codons (table S5-S6). Only one codon, CUC, coding for leucine, was overrepresented in all species of the dataset, with minimal RSCU values > 1.6. Consistent with overrepresented codons in the dataset ending with cytosine, GC3 percentages were higher than 50% for all species, and ranged from 54.1% in *Podospora fimiseda* (*Podosporaceae*) to 80.7% in *Crassicarpon thermophilum* (*Chaetomiaceae*) (table S6). Three codons (AUA, UUA and GUA, encoding isoleucine, leucine and valine, respectively) were underrepresented in all species in the dataset with maximum RSCU values < 0.6. All these three codons contained adenine (A) at the third codon position.

**Figure 2:**
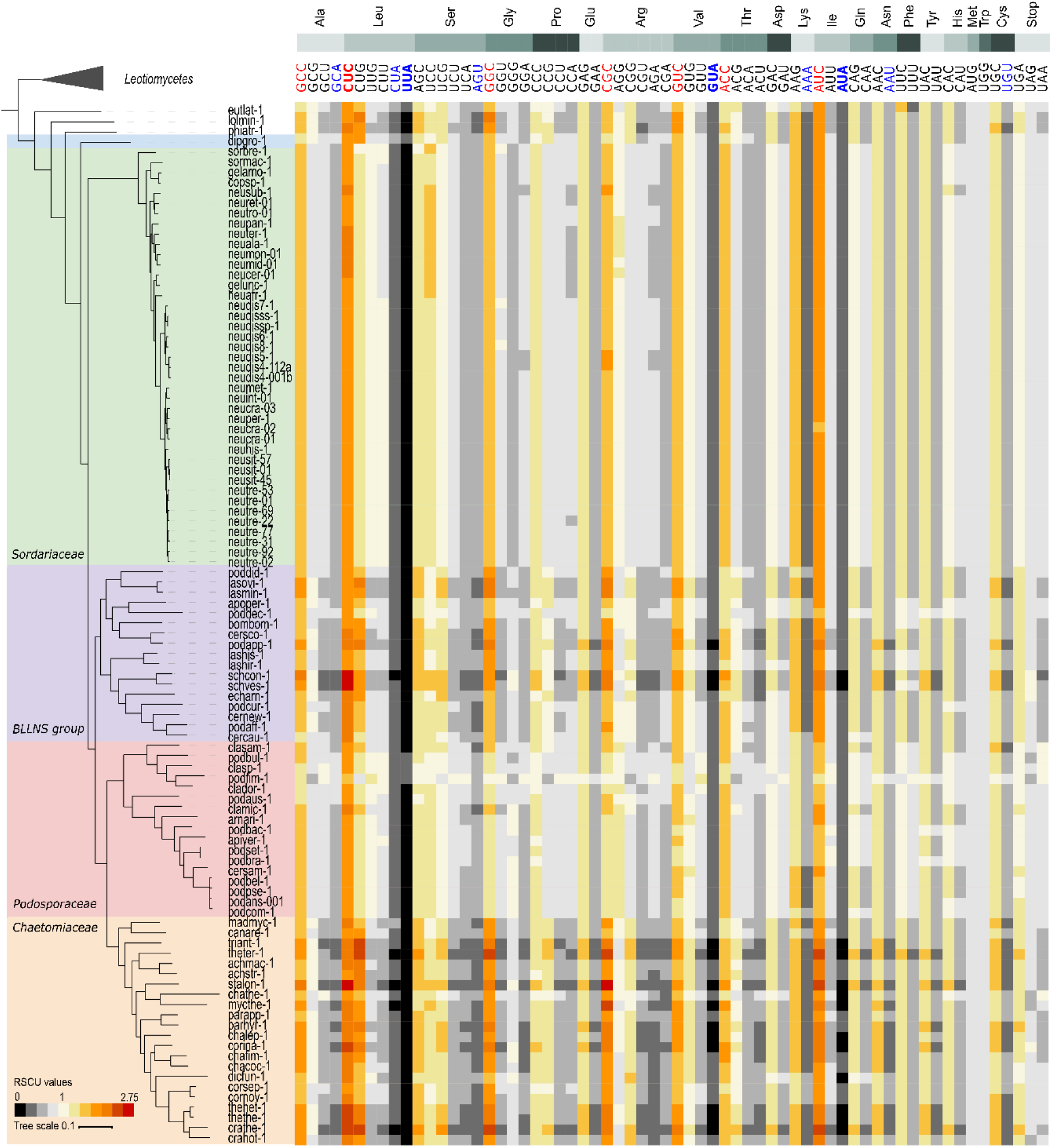
Patterns of relative synonymous codon usage across Sordariales (RSCU). Individual codons for each amino acid, ordered from most used to least used for the average of all genomes combined. RSCU values > 1 indicate that there is a higher frequency of a particular codon in the genome than expected under random use, while RSCU values <1 indicate that a codon is less frequent within the genome. Codons with RSCU values >1.6 or <0.6 are seen as over- and underrepresented codons. Heatmap colors range from black-grey (RSCU <1), to white (RSCU = 1), to red-darkred (RSCU>1). Codons with average or minimal RSCU > 1.6 are indicated with red letters, or red and bold letters, respectively. Codons with average or maximal RSCU < 0.6 are indicated with blue letters, or blue and bold letters, respectively.

In contrast to amino acid composition, which was confirmed in this study to remain largely similar across species and kingdoms, preferential usage of codons often differs between groups of organisms. Within the plant kingdom, for example, grasses and Caryophyllaceae plants preferred GC ending codons, while cruciferous plants, Rosaceae, Leguminosae and Pinaceae preferred AU ending codons (55). Within the kingdom fungi, GC ending codons are underrepresented within the Saccharomycotina, including in most budding yeasts (56,57), but overrepresented in Taphrinomycotina, and Pezizomycotina (57). The current study showed that GC3 overrepresentation is also found in all studied Sordariales genomes, but levels of overrepresentation are variable amongst groups.

### 3.3 Variation in codon usage bias across the Sordariales

Despite similar trends in codon usage across the order Sordariales, the four groups in the dataset (*Chaetomiaceae, Podosporaceae, Sordariaceae* and BLLNS) showed differences in the level of over- and underrepresentation (figure 2; table S5). Specifically, the *Chaetomiaceae* and the BLLNS showed a higher overall bias than the other two groups. Consistent with the higher overall CUB in these two groups, and order-wide overrepresentation of G and C ending codons, *Chaetomiaceae* and the BLLNS had the highest levels of GC3, with a mean of 73% and 69% respectively. Both *Chaetomiaceae* and the BLLNS showed significantly higher GC3 levels than *Podosporaceae* and *Sordariaceae* (p < 0.001).

The *Podosporaceae* showed the least bias of the four groups, with overall RSCU values closer to 1.0 than the *Sordariaceae, Chaetomiaceae* and the BLLNS. The *Podosporaceae* had the lowest GC3 content, with an average GC3 of 63%. Within the *Podosporaceae, Podospora fimiseda*, in particular, stood out with lower overall bias. *P. fimiseda* showed more RSCU values closer to 1 than its sister species, or than other genomes in the dataset. In contrast to the trends for the entire dataset, the low frequency codons in *P. fimiseda* were all G or C ending, while higher frequency codons in this specific species all contained A or U at the third codon position. Consistent with this finding, *P. fimiseda* had the lowest overall GC3 content of the dataset, with a GC3 content of 54%.

Differences in GC3 between the *Podosporaceae* and *Sordariaceae* were small (p = 0.47). The *Sordariaceae* showed slightly higher levels of bias than the *Podosporaceae*, with an average GC3 of 64%. Overall, less intra-family variation was seen within the *Sordariaceae* compared to other families. To this end, we note that the genome sequences of the taxa sampled are unevenly distributed across the groups of Sordariales. Specifically, within *Sordariaceae*, several closely related *Neurospora* species are over-represented compared to other taxa (21), which could explain high levels of RSCU similarity found within the *Sordariaceae* family.

### 3.4 Natural selection is the main driver for codon usage bias in the Sordariales

The effective number of codons (ENC) ranged from 41.6 in the *Crassicarpon thermophilum* genome (*Chaetomiacaeae*), to 57.3 in the genome of *Podospora fimiseda* (*Podosporaceae*). As ENC theoretically ranges from 20 (for a single codon per amino acid) to 61 (when all codons are used equally), our data indicated that the order harbors genomes with medium to almost no CUB on the genome-wide level. All genomes showed large differences in CUB between highly expressed genes and the rest of the genomes, as interpreted by having a codon adaptation index (CAI) CAI < 0.59 (26). Together, the low to medium overall levels of ENC and CAI suggest low to intermediate codon bias for the whole genome data in the investigated group of fungi (24), with large to intermediate difference in CUB between highly expressed genes and the rest of the genome (26).

Genome-wide CUB patterns were mainly formed by selection, indicated by shallow slopes in the neutrality plots. Significant negative correlations were found between GC12 and GC3 (R < 0, p < 0.01) in the majority of genomes (figure 3; table S7). The maximum absolute slope value of 0.31 showed that genome-wide natural selection driving codon usage was > 69% in all species in the dataset. In four genomes, the correlation between GC12 and GC3 was non-significant, also indicating that natural selection was the dominant driving force in shaping CUB patterns (43–45)(table S7).

**Figure 3:**
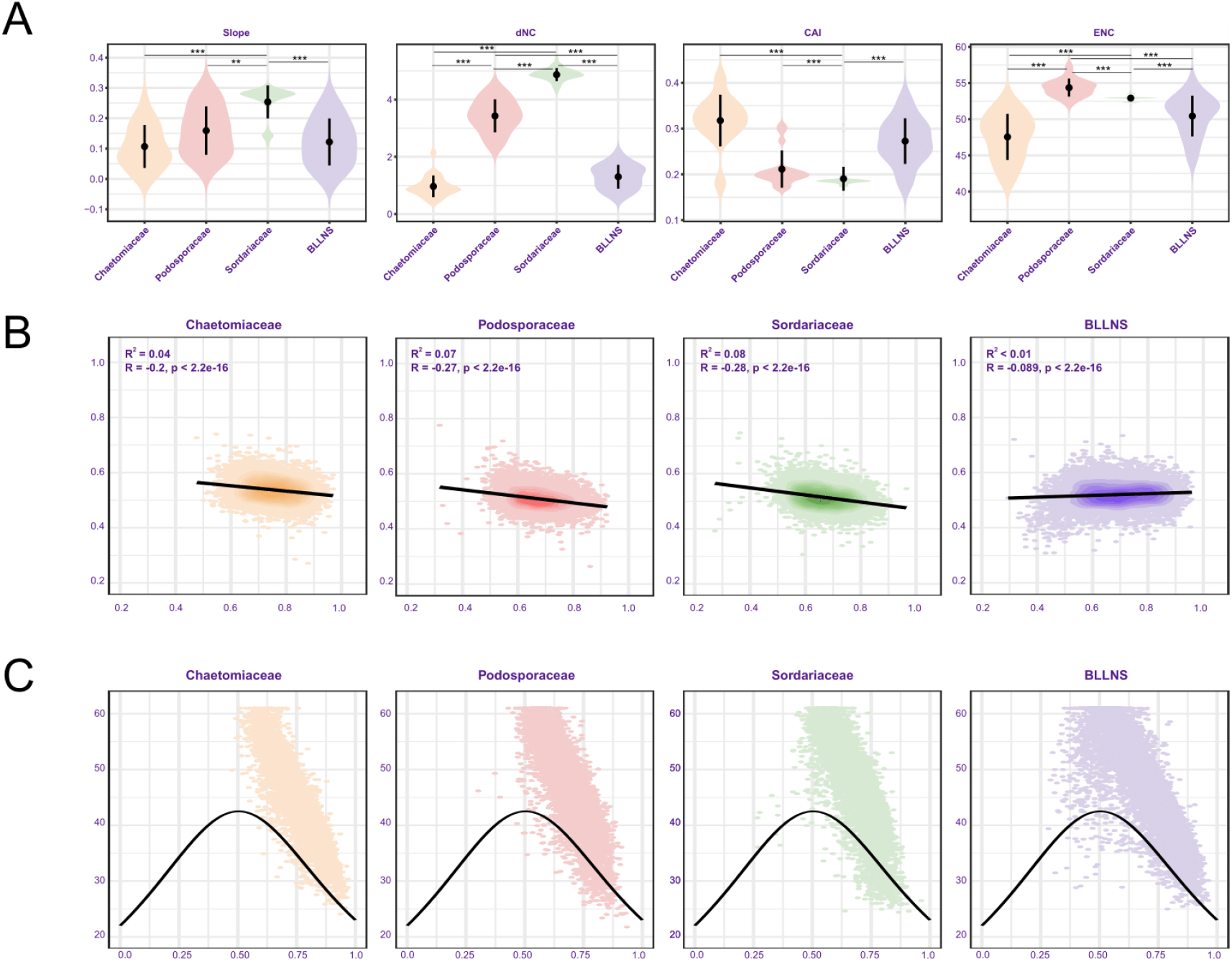
patterns of codon usage bias across Sordariales. (A) Violin plots for codon usage bias indices. From left to right: absolute slope of neutrality plots, dNC, CAI and ENC for *Chaetomiaceae, Podosporaceae, Sordariaceae*, and the BLLNS. Significance is shown for * p < 0.05, ** p < 0.01, *** p < 0.001 (B) Representative neutrality plots (GC12/GC3 plots) from each of the main groups. From left to right: *Chaetomiaceae (Chaetomidium leptoderma), Podosporaceae (Arnium arzonense), Sordariaceae (Neurospora sitophila-1)*, BLLNS group *(Bombardia bombarda)*. (C) Representative ENC/GC3 plots from each of the main groups, using the same genomes as B.

Finally, we created ENC/GC3 curves to reveal the relationship between nucleotide composition and codon bias in single genes. All genomes showed similar trends of ENC/GC3 correlations (figure 3). Placement of the majority of genes far above the expected ENC/GC3 curve indicated low levels of selection on usage of specific codons for the majority of genes in the dataset (24). The percentage of genes under selection for codon usage (dNC) ranged from 0.4% in *Achaetomium strumarium* (*Chaetomiaceae*) to over 4% for all *Sordariaceae* species, with a maximum of 5.4% for *Sordaria brevicollis*.

#### 3.4.1 *Chaetomiaceae* and the BLLNS contain the highest genome-wide levels of selection driving codon usage

Many genomes in the Sordariales showed similar trends of CUB and selection, but levels of CUB and the role of selection in shaping codon usage varied across the groups (figure 3). The highest levels of CUB and selection were found in the *Chaetomiaceae*, closely followed by the BLLNS group. *Podosporaceae* contained lower levels of CUB, and closely resembled *Sordariaceae*, which contained the lowest genome-wide CUB. *Chaetomiaceae, Podosporaceae*, and *Sordariaceae* were significantly different in ENC, CAI, and dNC. The BLLNS closely resembled the *Chaetomiaceae*, and no significant differences were found between the two for any of the analyzed variables (figure 3).

*Chaetomiaceae* showed the highest levels of CUB at the whole genome level. Amongst the *Podosporaceae, Sordariaceae* and *Chaetomiaceae* families, the *Chaetomiaceae* had significantly the lowest ENC values (average ENC = 47.5), and the highest CAI. The average CAI of 0.32 indicated large CUB differences between highly expressed genes and genome-wide CDS (26), but CAI was significantly lower than in *Sordariaceae* and *Podosporaceae. Chaetomiaceae* displayed the smallest dNC of the four groups, 1.0% of genes on average. Thus, consistent with the relatively high CAI, very few genes showed increased levels of selection compared to genome-wide trends. The average absolute slope of neutrality plots was closest to zero, with an average absolute slope of 0.11, indicating 89% selective constraint on genome-wide codon usage.

The BLLNS group showed similar trends of CUB as the *Chaetomiaceae*, indicated by the relatively low ENC (average = 50.4), and relatively high CAI (average = 0.27). The average selection level was 88%, indicated by an average absolute slope in the neutrality plots of 0.12. Similar to the *Chaetomiaceae*, the relatively high CAI is accompanied by a low percentage of genes that showed selection as driver of codon usage, with an average dNC of 1.3%. Together, the results indicated high CUB and selection driving codon usage in the BLLNS, but slightly lower than in the *Chaetomiaceae*. Levels of codon usage bias were higher in the BLLNS than in the *Podosporaceae*; The BLLNS had a significantly lower percentage of genes under selection and lower ENC values than the *Podosporaceae*, but no significant differences were found for the average CAI and the absolute slope of the neutrality plot (figure 3).

The *Podosporaceae* showed low to intermediate levels of CUB. The CAI in *Podosporaceae* genomes (average CAI = 0.21) is significantly lower than in the *Chaetomiaceae*, and significantly higher than in *Sordariaceae* genomes. The ENC was significantly highest in the *Podosporaceae* family (average ENC = 54.4), indicating low pressure to use certain codons over others. No significant difference was found for the absolute slope of the neutrality plots between *Podosporaceae* and *Chaetomiaceae* or the BLLNS. The average absolute slope remained low, at 0.16, indicating that codon usage was driven by selection in more than >80% of the genome. The *Podosporaceae* contained an intermediate level of genes under selection driving codon usage, on average 3.4%. The *Podosporaceae* genomes contained slightly higher levels of codon bias and selection than *Sordariaceae* genomes, but overall differences between the two families were small; CAI, dNC and slope of neutrality plots were significantly smaller in *Podosporaceae* compared to *Sordariaceae*, while ENC was significantly higher in the *Podosporaceae*.

The *Sordariaceae* showed the lowest levels of CUB and the lowest levels of selection on genome-wide codon usage. *Sordariaceae* contained a high ENC (average ENC = 52.9), indicative of low codon bias. They contained significantly the lowest levels of CAI, 0.19 on average. As such, *Sordariaceae* genomes contained the lowest genome-wide levels of codon usage bias, and the largest differences in CUB between highly expressed genes and the rest of the genome. *Sordariaceae* genomes further contained relatively more genes that showed codon usage driven by selection (average dNC = 4.9%). Consistent with the low genome-wide codon bias, the *Sordariaceae* genomes showed a relatively large impact of mutational bias relative to selection. The average absolute slope of the neutrality plot was 0.25, indicative of codon usage being driven for 25% by mutational bias versus 75% selection.

Overall, genomes from the Sordariales showed low to intermediate genome-wide codon usage bias, with large to intermediate difference in CUB between highly expressed genes and the rest of the genome. Genome-wide CUB patterns were mainly formed by selection, and levels of bias and selection were dependent on the researched groups. Codon bias was found to be highest in the *Chaetomiaceae*, followed by the BLLNS, *Podosporaceae* and lastly *Sordariaceae*, which contained the lowest levels of genome-wide codon bias.

#### 3.4.2 Signs of selection driving codon usage in *Sordariaceae* driven by a small subset of genes

Our results indicated that, for the different genomes, higher percentages of genes under selection driving codon usage (increased dNC) corresponded to lower genome-wide codon usage (increased ENC, decreased CAI) and lower selection on genome-wide codon usage (increased slope of neutrality plots) (figure 3). Indeed, even after accounting for phylogeny, pearson’s correlation coefficient showed that higher percentages of genes under selection were significantly correlated with lower genome-wide codon usage bias, as indicated by strong positive correlations between ENC and dNC (R = 0.51, p < 0.001), and negative correlations between CAI and dNC (R = -0.46, p < 0.001). Higher percentages of genes where codon usage is driven by selection were furthermore correlated with lower genome-wide selection driving codon usage, with a positive correlation between dNC and the absolute slope of neutrality plots (R=0.46, p < 0.001). Thus, the stronger the genome-wide codon usage bias and selection that drives genome-wide codon usage, the smaller the percentage of genes showing signs of selection.

The negative correlation between dNC and CAI further indicated that the difference in CUB between highly expressed genes and the rest of the genome increased as the percentage of genes under selection for codon usage increased. We further investigated these correlations in Sordariales genomes by analyzing patterns of RSCU values across genes that showed signs of selection (i.e. those below the ENC/GC3 curve). As expected by the low order-wide CAI values, the subset of genes that showed high levels of selection driving codon usage showed higher CUB compared to the whole-genome CDS (figure 4): RSCU values generally deviated further from 1 than in genome-wide analysis (figure 2, figure 4). For all genomes, GC-ending codons were preferred over AT ending codons. Average GC3 levels were higher in this group of genes compared to genome-wide CDS, and ranged from 67% in *Lasiosphaeria miniova* (BLLNS group) to 83% in *Staphylotrichum longicolleum* (*Chaetomiaceae*).

**Figure 4:**
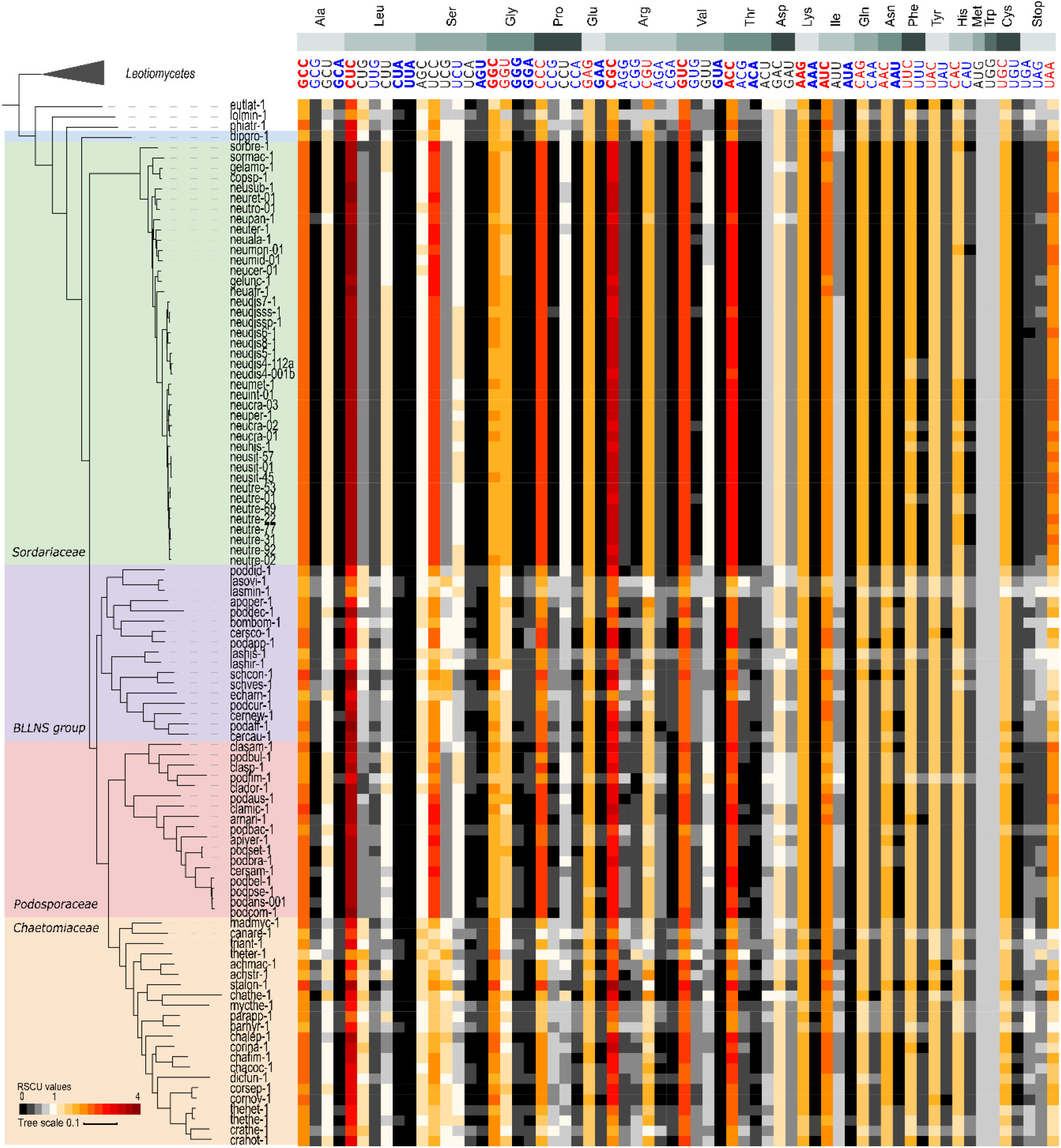
Patterns of relative synonymous codon usage for genes under selection. Amino acids are ordered in the same order as figure 1, with individual codons ordered as in figure 2. RSCU values larger than 1 indicate that there is a higher frequency of a particular codon in the genome than expected under random use, while RSCU values <1 indicate that a codon is less frequent within the genome. Codons with RSCU values >1.6 or <0.6 are seen as over- and underrepresented codons. Heatmap colors range from black-grey (RSCU <1), to white (RSCU = 1), to red-darkred (RSCU>1). Codons with average or minimal RSCU > 1.6 are indicated with red letters, or red and bold letters, respectively. Codons with average or maximal RSCU < 0.6 are indicated with blue letters, or blue and bold letters, respectively.

Lower CAI values suggest that selection acts only on highly expressed genes, and that there are larger differences in CUB between highly expressed genes and the rest of the genome (26). Indeed, CUB trends in this subset of genes were opposite to whole genome trends. In genome-wide data, *Chaetomiaceae* showed the highest bias, followed by BLLNS, while *Podosporaceae*, and *Sordariaceae* contained the least bias. In genes showing signs of selection, *Sordariaceae* showed the highest bias, with RSCU values deviating the furthest from 1 (figure 4). The GC3 was significantly highest in the *Sordariaceae* (average = 76%, p <0.01), compared to all other groups. Furthermore, only the *Sordariaceae* showed a significant difference in GC3 content of genome-wide CDS (average GC3 = 64%) versus GC3 content of genes under selection (p < 0.05).

The *Sordariaceae* was closely followed by the *Podosporaceae*, which contained slightly lower levels of codon bias in this subset of genes (figure 4). *Chaetomiaceae* and the BLLNS showed the least bias, with many RSCU values closer to 1, and relatively low GC3 differences between genome-wide data and this subset of genes. We found no significant GC3 differences amongst the *Podosporaceae* (average GC3 = 73.6%), *Chaetomiaceae* (average GC3 = 74.2%), and the BLLNS (average GC3 = 73.2%). However, the *Podosporaceae* contained larger differences in genome-wide GC3 (average GC3 = 63%) versus GC3 in the subset of genes under selection (average GC3 = 73.6%). These differences were much smaller in *Chaetomiaceae* (average genome-wide GC3 = 73%) and the BLLNS (average genome-wide GC3 = 69%). As such, we expect that the *Chaetomiaceae* and BLLNS experienced higher selection on codon usage throughout the entire genome, rather than selection driven by a small number of genes.

### 3.5 Patterns of codon usage in *Sordariaceae* and *Podosporaceae* resemble the ancestral state of Sordariales

The Sordariales phylogeny places *Chaetomiaceae* and *Podosporaceae* as sister clades, while *Sordariaceae* clusters in a separate clade away from both of these families and the BLLNS (figure 1)(21). Visual inspection of figure 1-4 showed similarity in amino acid composition, patterns of CUB, and overall strength of selection driving codon usage between the *Chaetomiaceae* and the BLLNS. Traits in the *Podosporaceae*, however, more closely resembled those of the *Sordariaceae*. To more systematically investigate the similarity across the order, we used automatic clustering of the genomes based on both amino acid composition and RSCU values (figure S1 and S2, respectively). For both indices, the automatic clustering of genomes confirmed that *Sordariaceae* and *Podosporaceae* were more similar to each other than each of them was to any of the other clades, even though they are not sister-groups in the phylogeny.

One factor that could explain the similarities found between *Chaetomiaceae* and the BLLNS could be that the grouping of families in the phylogeny is incorrect. However, the phylogeny is based on genome-wide information and shows good support (21). Additionally, the trends seen for CUB and amino acid composition do not hold true for some other researched genomic traits. For example, the average repeat percentage of *Podosporaceae* is lower than that of either the *Chaetomiaceae* or *Sordariaceae* (21). We therefore find it likely that the family branching in the phylogenetic tree is correct.

We next reconstructed ancestral states of GC3 in a wide range of fungi (GC3 values and phylogeny obtained from Wint et al., 2022). The Wint dataset contains seven Sordariales species, of which five *Chaetomiaceae* and two *Podosporaceae* genomes. Similar to results presented here, the GC3 values of the *Chaetomiaceae* were higher than those of the *Podosporaceae* (table S8). Our ancestral state reconstruction (figure S3), showed that the Sordariales contain a relatively high GC3 content, but that the GC3 content decreases consistently towards the root of the Sordariomycetes. Based on these analyses, we hypothesize that the GC3 values, amino acid composition, and CUB of both *Podosporaceae* and *Sordariaceae* represent the ancestral state of Sordariales. The other two lineages – that of *Chaetomiaceae* and the BLLNS, show changes from that state, with an increase in natural selection driving use of specific codons, and higher genome-wide CUB (figure S1 and S2).

### 3.6 Thermophilia and increased codon bias in Chaetomiaceae

The *Chaetomiaceae* and the BLLNS showed changes from the ancestral state of the Sordariales with increase in natural selection driving use of specific codons. Based on a number of previous reports, it can be hypothesized that increased CUB, at least in *Chaetomiaceae*, is associated with ecological niche specialization. Botzman and Margalit (2011) found that prokaryotes living in specialized habitats – including those living in high temperatures - showed significantly lower ENC and higher CAI, indicating stronger selection on genome-wide codon usage compared to organisms living in multiple habitats (26). Trends seen in both the BLLNS and the *Chaetomiaceae* are comparable to those in specialized prokaryotes. It is impossible at this stage to speculate on the cause of divergence within the BLLNS, as our dataset contains only a few species divided over five different families. In the *Chaetomiaceae*, however, increased codon usage bias has previously been correlated with fungal thermophilia (18). Optimal growth temperature has been described as a major factor affecting codon usage in prokaryotes (58), and fungal thermophiles have previously been described to show increased GC3 content, and decreased ENC, compared to closely related non-thermophilic fungi (18). Our study confirmed that *Chaetomiaceae* had the highest levels of genome-wide CUB and genome-wide selection of the Sordariales, and, similar to other specialist fungal species, *Chaetomiaceae* lack large differences in CUB between highly expressed genes and other genes in the genome (59).

We note, however, that thermophilic lifestyle cannot currently be established as the sole driver of the high codon usage bias in this group of fungi. First of all, GC3 levels are not always increased in thermophilic *Chaetomiaceae* fungi (18). Within our study, codon bias levels largely overlap amongst genomes belonging to *Chaetomiaceae*, and no significant differences in CUB indices were found amongst species with different optimal growth temperatures (table S9, figure S4). Our results further show discrepancies with previously described trends in amino acid frequencies. Three fungal thermophiles investigated by van Noort and colleagues (2013) showed increased levels of arginine and tryptophane when compared to closely related fungi with lower optimal growth temperatures. These trends were not visible in our dataset, and both tryptophan and arginine frequencies did not significantly differ amongst *Chaetomiaceae* with different optimal growth temperatures (table S9, figure S4). To this end, it should be mentioned that it is difficult to be certain about the classification of optimal growth temperatures. Classification of thermophilia differs between different studies, and different studies describe slightly different optimal growth temperatures for the same fungal strains present in our dataset (18,19). The reason for the discrepancies in optimal growth temperatures, amino acid frequencies and codon bias trends between studies is currently not clear, and more research is needed to further detangle the correlations amongst optimal growth temperature, genomic traits, and CUB trends in *Chaetomiaceae* fungi.

Overall, we expect that *Chaetomiaceae* have experienced ecological niche specialization with correlated increased selection pressure on use of certain codons over others, when compared to *Sordariaceae* and *Podosporaceae*, but it remains currently unclear whether this is caused by high temperature environments or other types of niche specialization. In contrast to highly specialized *Chaetomiaceae*, the ancestral state of the Sordariales could be indicative of a more generalist nature. Prokaryotic species living in less specialized conditions, and a wide range of habitats, show low genome-wide CUB, but high levels of CUB in a subset of genes (26,59,61). For example, Badet et al. (2017) showed that generalist parasitic fungi contain a set of genes with high CUB, compared to genome-wide CDS with lower biases. The same trends were found in the majority of *Sordariaceae* and *Podosporaceae* genomes analyzed in this project. Although a generalist nature is harder to define than ecological specialization, *Neurospora* species have been described to generally lack host specificity and to have broad geographic ranges (62). Over the entire dataset, the ecological niche of many species remains unclear (see e.g. 15,63), but the low genome-wide CUB and the large differences in CUB between highly expressed genes and the rest of the genome would be in line with a more generalist nature than that seen in *Chaetomiaceae*.

## Data and materials

The computations/data handling were enabled by resources provided by the Swedish National Infrastructure for Computing (SNIC) at UPPMAX and DARDELL, partially funded by the Swedish Research Council through grant agreement no. 2018-05973.

