## Supplementary figures for "Patterns and drivers of genome-wide codon usage bias in the fungal order Sordariales"

Figure s1

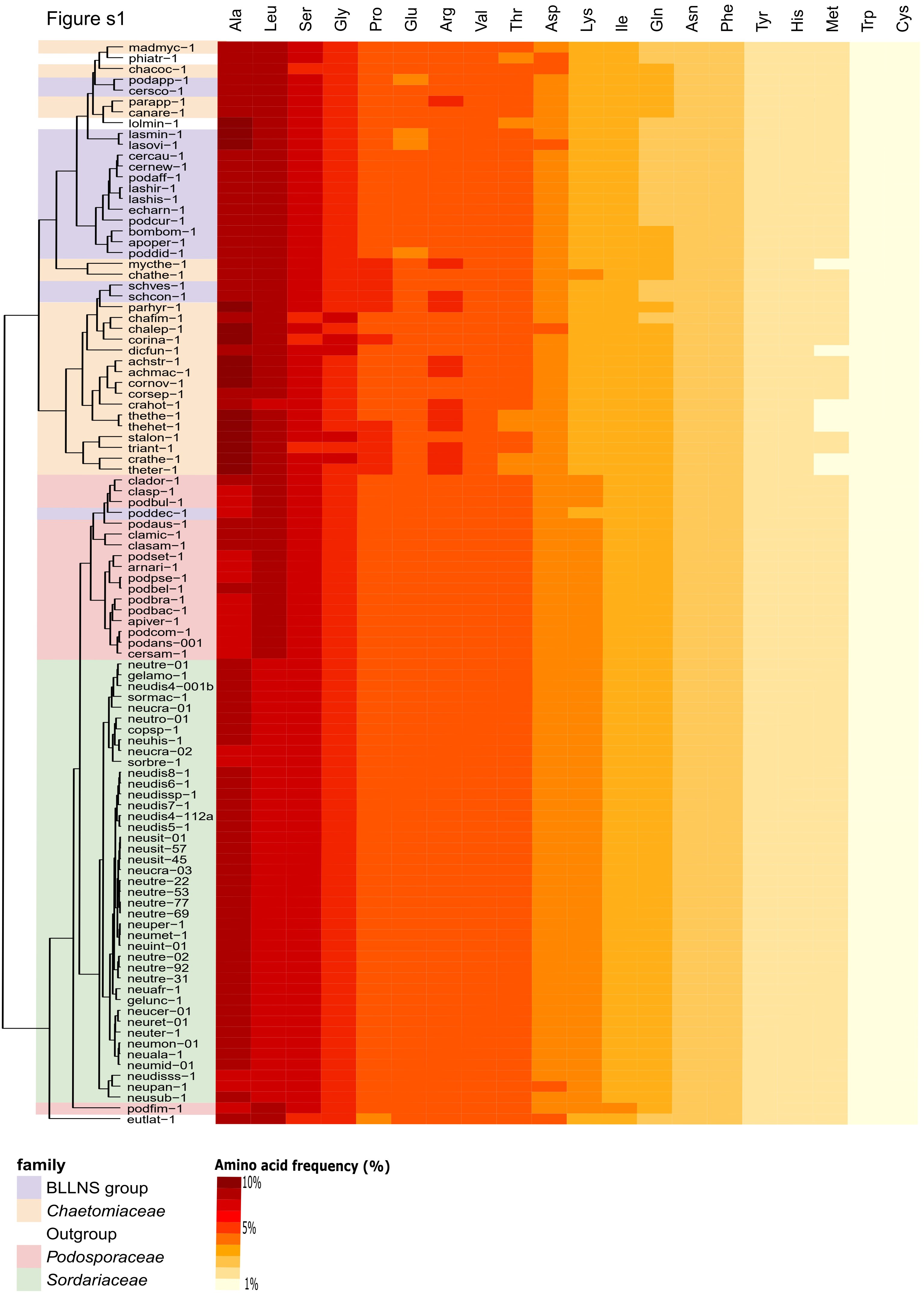

Ala
Leu
Ser
Gly
Pro
Glu
Arg
Val
Thr
Asp
Lys
Ile
Gln
Asn
Phe
Tyr
His
Met
Trp
Cys
madmyc−1
phiatr−1
chacoc−1
podapp−1
cersco−1
parapp−1
canare−1
lolmin−1
lasmin−1
lasovi−1
cercau−1
cernew−1
podaff−1
lashir−1
lashis−1
echarn−1
podcur−1
bombom−1
apoper−1
poddid−1
mycthe−1
chathe−1
schves−1
schcon−1
parhyr−1
chafim−1
chalep−1
corina−1
dicfun−1
achstr−1
achmac−1
cornov−1
corsep−1
crahot−1
thethe−1
thehet−1
stalon−1
triant−1
crathe−1
theter−1
clador−1
clasp−1
podbul−1
poddec−1
podaus−1
clamic−1
clasam−1
podset−1
arnari−1
podpse−1
podbel−1
podbra−1
podbac−1
apiver−1
podcom−1
podans−001
cersam−1
neutre−01
gelamo−1
neudis4−001b
sormac−1
neucra−01
neutro−01
copsp−1
neuhis−1
neucra−02
sorbre−1
neudis8−1
neudis6−1
neudissp−1
neudis7−1
neudis4−112a
neudis5−1
neusit−01
neusit−57
neusit−45
neucra−03
neutre−22
neutre−53
neutre−77
neutre−69
neuper−1
neumet−1
neuint−01
neutre−02
neutre−92
neutre−31
neuafr−1
gelunc−1
neucer−01
neuret−01
neuter−1
neumon−01
neuala−1
neumid−01
neudisss−1
neupan−1
neusub−1
podfim−1
eutlat−1
family
BLLNS group
Chaetomiaceae
Outgroup
Podosporaceae
Sordariaceae
Amino acid frequency (%)
10%
5%
1%

Figure s1: Amino acid composition across the Sordariales, with automatic clustering of genomes. Average amino acid frequencies (%) for all genomes. Amino acids are ordered from most used (left, dark-red) to least used (right, light-yellow). *Sordariaceae* and *Podosporaceae* cluster together, and *Chaetomiaceae* and the BLLNS group cluster together.

Figure s2

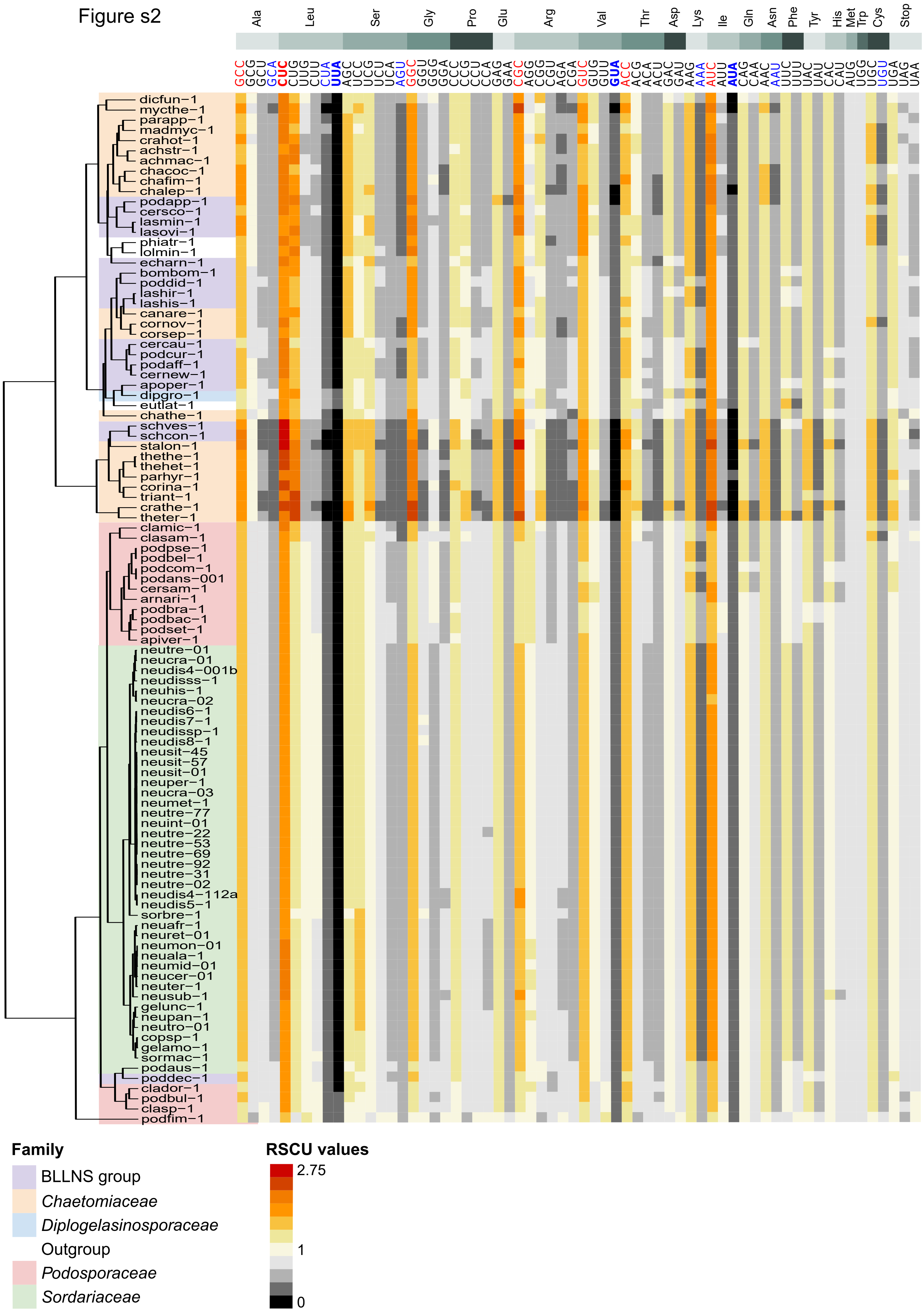

Ala
Leu
Ser
Gly
Pro
Glu
Arg
Val
Thr
Asp
Lys
Ile
Gln
Asn
Phe
Tyr
His
Met
Trp
Cys
Stop
GCC GCG GCU GCA CUC CUG UUG CUU CUA UUA AGC UCC UCG UCU UCA AGU GGC GGU GGG GGA CCC CCG CCU CCA GAG GAA CGC AGG CGG CGU AGA CGA GUC GUG GUU GUA ACC ACG ACA ACU GAC GAU AAG AAA AUC AUU AUA CAG CAA AAC AAU UUC UUU UAC UAU CAC CAU AUG UGG UGC UGU UGA UAG UAA
dicfun−1
mycthe−1
parapp−1
madmyc−1
crahot−1
achstr−1
achmac−1
chacoc−1
chafim−1
chalep−1
podapp−1
cersco−1
lasmin−1
lasovi−1
phiatr−1
lolmin−1
echarn−1
bombom−1
poddid−1
lashir−1
lashis−1
canare−1
cornov−1
corsep−1
cercau−1
podcur−1
podaff−1
cernew−1
apoper−1
dipgro−1
eutlat−1
chathe−1
schves−1
schcon−1
stalon−1
thethe−1
thehet−1
parhyr−1
corina−1
triant−1
crathe−1
theter−1
clamic−1
clasam−1
podpse−1
podbel−1
podcom−1
podans−001
cersam−1
arnari−1
podbra−1
podbac−1
podset−1
apiver−1
neutre−01
neucra−01
neudis4−001b
neudisss−1
neuhis−1
neucra−02
neudis6−1
neudis7−1
neudissp−1
neudis8−1
neusit−45
neusit−57
neusit−01
neuper−1
neucra−03
neumet−1
neutre−77
neuint−01
neutre−22
neutre−53
neutre−69
neutre−92
neutre−31
neutre−02
neudis4−112a
neudis5−1
sorbre−1
neuafr−1
neuret−01
neumon−01
neuala−1
neumid−01
neucer−01
neuter−1
neusub−1
gelunc−1
neupan−1
neutro−01
copsp−1
gelamo−1
sormac−1
podaus−1
poddec−1
clador−1
podbul−1
clasp−1
podfim−1
Family
BLLNS group
Chaetomiaceae
Diplogelasinosporaceae
Outgroup
Podosporaceae
Sordariaceae
RSCU values
2.75
1
0

Figure s2: Patterns of relative synonymous codon usage, with automatic clustering of genomes. Amino acids are ordered in the same order as figure 1, with individual codons ordered as in figure 2. RSCU values larger than 1 indicate that there is a higher frequency of a particular codon in the genome than expected under random use, while RSCU values <1 indicate that a codon is less frequent within the genome. Codons with RSCU values >1.6 or <0.6 are seen as over- and underrepresented codons. Heatmap colors range from black-grey (RSCU <1), to white (RSCU = 1), to red-darkred (RSCU>1).

Figure s3

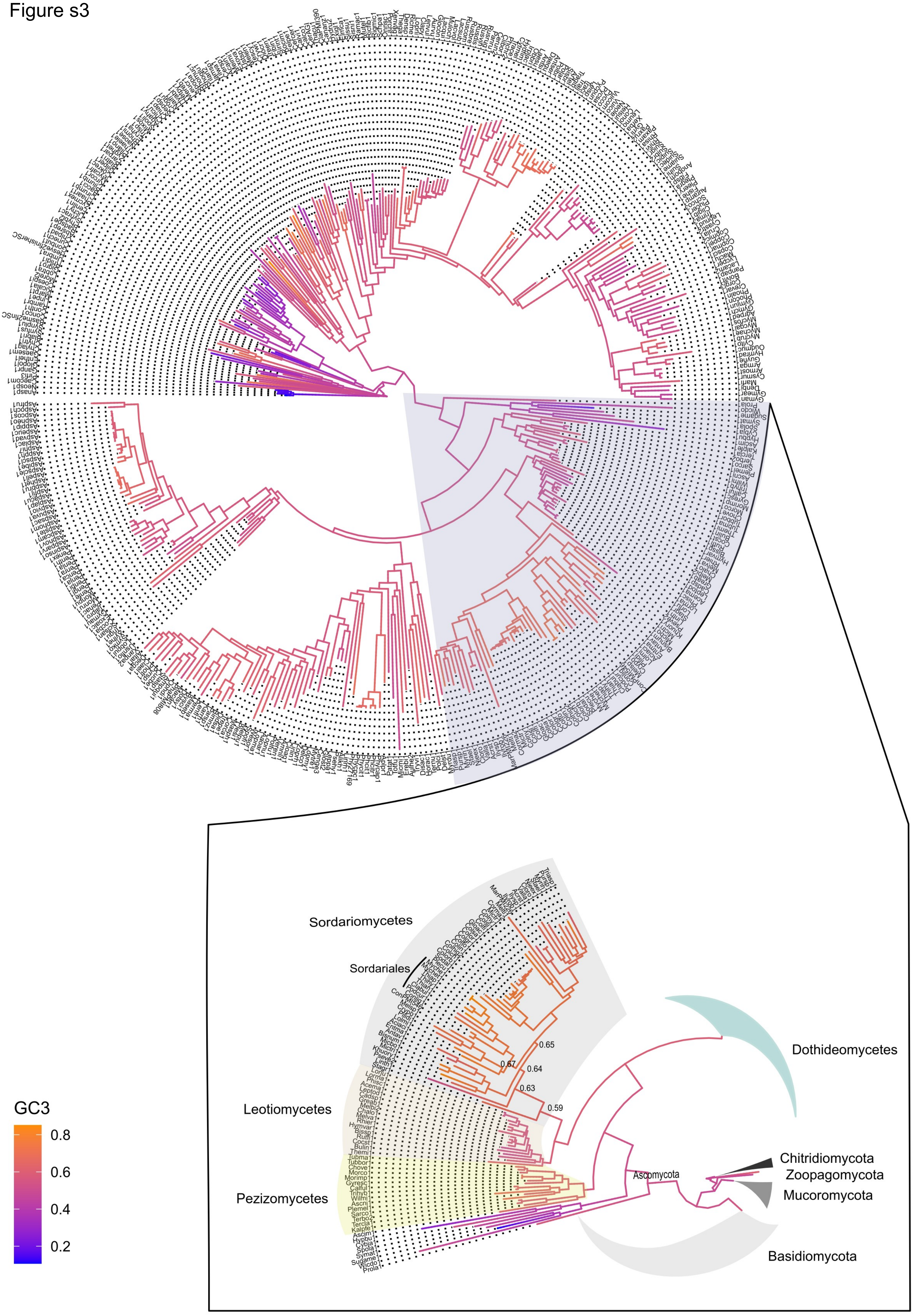

Sordariomycetes
Sordariales
Leotiomycetes
Pezizomycetes
Dothideomycetes
Ascomycota
Chitridiomycota
Zoopagomycota
Mucoromycota
Basidiomycota
0.65
0.67
0.64
0.63
0.59
GC3
0.8
0.6
0.4
0.2

Figure s3: Ancestral state reconstruction of GC3 values across the fungal kingdom. GC3 values were obtained from Wint et al., 2022 and reconstructed on the species phylogeny. The heatmap denotes ancestral GC3 frequencies from small (dark blue) to high (orange). Ancestral states are shown from the most recent common ancestor of the Sordariomycetes to the root of the Sordariales order.

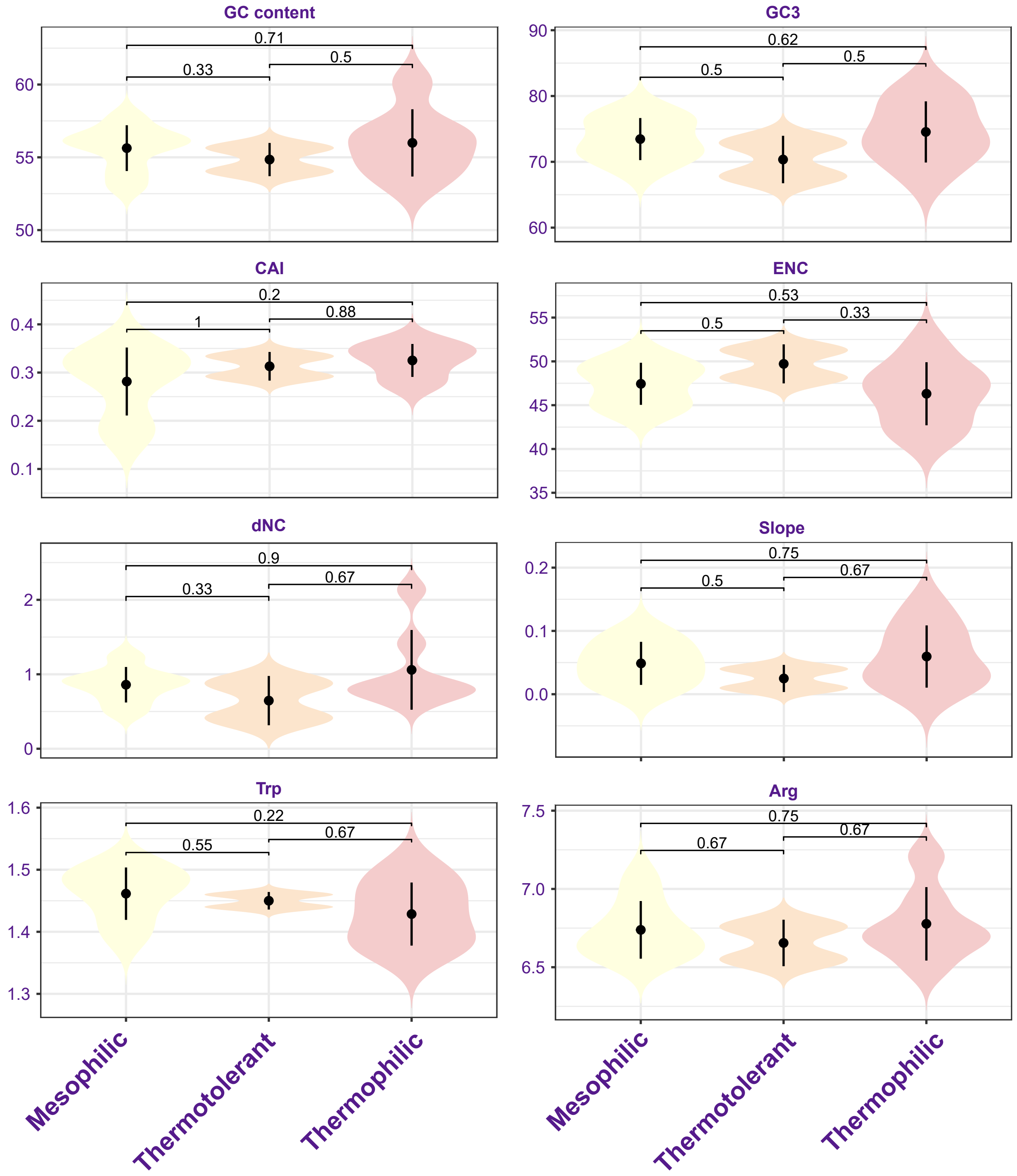


Figure s4: patterns of codon usage bias across Chaetomiaceae species with different optimal growth temperatures. Optimal growth temperatures obtained from Steindorf et al., 2024 and van den Brink et al., 2015. From left to right for mesophilic species (optimal growth temperature < 37*C), thermotolerant species (optimal growth temperature between 37-45*C), and thermophilic species (optimal growth temperature at or above 45*C). P-values were obtained with an unpaired wilcoxon rank sum test.
